# Irrational groups: decoy placement, not group size, shapes collective shoal-size preference in adult zebrafish

**DOI:** 10.64898/2026.02.09.704858

**Authors:** Abhishek Singh, Nandini Bhattacharya, Mrittwika Dutta Gupta, Thanvi Sajith, Manavi Gupta, KR Srinandini, Bittu Kaveri Rajaraman

## Abstract

Group living often improves decision accuracy in animals, yet whether increasing group size buffers against context-dependent biases remains unexplored. One such bias is the decoy effect, where the presence of a third option shifts preferences between two alternatives. Here, we tested whether introducing a decoy shoal influences collective preference for the larger female shoal in adult male zebrafish (*Danio rerio*), and whether the strength of this effect depends on group size. Groups of two, three, or four males were presented with female shoals under two choice contexts: a dichotomous contrast (four vs. two fish) and a trichotomous contrast including an additional alternative of one, three, or five fish. The order of presentation (dichotomous-first or trichotomous-first) was counterbalanced, and multi-animal tracking was used to quantify group-level shoal-size preference (time spent near shoals), inter-individual distance (IID), polarization, and swimming speed. In the dichotomous-first order, across all group sizes, subject shoals consistently showed a baseline preference for the larger shoal. Adding a third decoy option altered this preference only when the decoy was extreme (one or five fish relative to the 4 vs. 2 alternatives), reducing relative preference toward indifference. Group IIDs were associated with context-dependent shifts in relative preference in the dichotomous-first order, whereas polarization and swimming speed were not. In the trichotomous-first order, groups showed no preference for the larger shoal, with or without a decoy shoal. Our results demonstrate that context-dependent biases, shape collective shoal choice, with effects driven by choice structure and order of presentation of options than by group size.

## Introduction

Improved decision-making is one of the key advantages of social living (Conradt & List 2009; Couzin 2009, Krause et al. 2010). Empirical studies show that groups frequently outperform solitary individuals in both the accuracy and speed of their decisions, with performance often improving further as group size increases (Ward et al. 2008; Sasaki & Pratt 2011). These benefits emerge through collective cognition, whereby individuals share, integrate, and refine information through social interactions (Couzin 2009). Across diverse taxa, groups benefit from shared information: fish (Ward et al. 2008), birds (Farine et al. 2014), mammals (Sueur et al. 2010; Gall et al. 2017), insect colonies (Prerna et al. 2012), amoeboid organisms (Reid & Latty 2016), and even human crowds (Faria et al. 2010) coordinate behavior by integrating socially acquired cues.

Decision-making in animals evolves under ecological and cognitive constraints, such that selection favours strategies that are fast and efficient rather than perfectly optimal (Houston 1997). These constraints often favor the evolution of simple heuristics that reduce computational costs by relying on limited information (Hutchinson & Gigerenzer 2005; Livnat & Pippenger 2008). Such heuristics may violate assumptions of economic rationality, which posit consistent, utility-maximising preferences (Rieskamp et al. 2006; Kacelnik 2006). These behaviors may still be biologically rational, in the sense that they may function effectively to increase fitness within their ecological, cognitive, and energetic constraints (Kacelnik 2006). One well-known example of a violation of economic rationality is the decoy effect, in which the presence of an additional option shifts preferences among alternatives (Huber et al. 1982, Hemingway et al. 2024).

A common explanation for improved accuracy in group decisions is that individual, randomly distributed errors counterbalance one another when many animals contribute their assessments (Surowiecki 2004). For instance, in migratory birds, each individual may hold an imprecise sense of direction, yet the collective integration of these noisy cues can yield a more precise group orientation (Simons 2004; Camazine 2020). However, simply aggregating individual errors cannot always account for systematic shifts in preference that arise in context effects. Collective decisions may involve a qualitative change in option evaluation, such that individuals do not assess alternatives independently; instead, comparisons can emerge from social interactions within the group, with animals relying more on social information than on solitary evaluation (Sasaki & Pratt 2011).

Nest-site choice studies in *Temnothorax* ants have demonstrated how collective decision-making can eliminate decoy-driven bias that is otherwise expressed at the individual level. Edwards and Pratt (2009) first challenged whole colonies with a classic asymmetric-dominance design: ants had to choose between two target nest sites that traded off entrance size and interior darkness, attributes known to influence nest quality, and then the same choices were repeated with the addition of a third nest asymmetrically dominated by one target but not the other. Colonies showed no shift in preference with either type of decoy, violating neither regularity nor the constant-ratio rule, even though the target nests were deliberately matched to make the decision difficult. Sasaki and Pratt (2011) expanded this framework by testing both individual ants and intact colonies using targets of closely matched attractiveness and two strategically placed decoys (one dominated by A, one by B). Individual ants, required to carry brood items into a chosen site, exhibited strong decoy effects; their preferences for A versus B were reversed simply by experiencing the corresponding decoy beforehand. In contrast, colonies given the same sets of nests, and induced to emigrate first into the decoy before choosing a final site, remained completely immune to the decoy’s influence. Robinson et al. (2014) took this further by removing the possibility of individual comparison altogether: by blocking ants from visiting both nests, they showed that colonies still made accurate, fast decisions. Demonstrating that collective choice in eusocial insects may not require individuals to directly compare alternatives and group-level evaluation emerges from distributed rules, recruitment, and quorum sensing.

In contrast to eusocial insect colonies—where high genetic relatedness, stable group membership, and specialised communication systems (such as pheromone trails or waggle dances) enable robust distributed decision-making—shoaling fish typically comprise groups of individuals that may or may not be related, have fluid shoal membership, and rely largely on passive cues derived from the position and movement of neighbours (Ioannou et al. 2011). Improved performance in larger fish shoals can often be explained by simpler mechanisms, including reduced individual vigilance, increased foraging motivation, or the influence of particularly competent individuals, rather than by genuinely distributed information integration (Sumpter & Pratt 2009; Ioannou et al. 2011; Ioannou 2017).

Whether groups of fish perform better than individuals on cognitive tasks has produced mixed results. In guppies, dyads outperform single individuals in numerical discrimination tasks (Bisazza et al. 2014), and groups show stronger preferences for vegetated over barren sector than individuals (Varracchio et al. 2025). The zebrafish dyads tested in a colour discrimination task do not outperform the more accurate member of the pair (Wang et al. 2015). The effect of group size on cognitive performance may also depend on ecological context. For example, in wild caught guppies, decision speed in high predation populations was unaffected by group size, whereas in low predation populations it peaked at intermediate group sizes and declined in the largest groups (Wade et al. 2020).

Fish shoals provide a tractable system for studying collective cognitive biases such as the decoy effect, owing to their loose social associations compared to eusocial insects. Yet it remains unexplored whether individual-level biases are expressed at the group level, and how their strength may be modulated by group size. In this study, we tested male zebrafish in groups of two, three, or four individuals using a shoaling preference assay in which they chose between female display shoals under a dichotomous context (four versus two fish) and a trichotomous context where a decoy shoal of one, three, or five fish was introduced. Using multi-animal tracking, we recorded the movements of each individual as the group performed the shoaling preference task, to model group-level preference for the larger shoal and extract measures of interindividual distance, polarization, and movement speed to assess how social decision making and group structure covary during the choice process. By comparing behaviour across group sizes and choice contexts, we test whether the decoy effect emerges at the group-level and whether its magnitude depends on group size. Based on evidence from eusocial insects indicating reduced context-dependent biases at the colony level, we predicted that the strength of context bias would decrease with increasing group size.

## Methods

### Subjects

Adult captive-bred zebrafish (Danio rerio; age < 1 year) were obtained from a local pet store in Daryaganj, New Delhi, India. The fish were housed in a ZebTec Active Blue Stand-Alone system (Tecniplast, PA, USA) at Ashoka University, Sonipat, Haryana, India. They were maintained under a 12:12 h light–dark cycle (lights on 10:00 a.m. to 10:00 p.m.) with water parameters kept at 28–30 °C, pH 7.5–8.5, and conductivity between 650–700 µS. Fish were fed ad libitum twice daily with powdered Tetra-Tetramin flakes.

Male zebrafish of comparable size were housed in groups of two, three, or four individuals in separate yet adjacent transparent 1.5 litre tanks within the ZebTec system. The arrangement allowed visual contact with neighboring groups. Fish were maintained under these group conditions for one week prior to the shoaling assays to allow social familiarity to develop. Adult females from the same population were randomly chosen as display fish for the shoal choice tests.

### Shoal choice apparatus

The shoaling apparatus consisted of a transparent cylindrical acrylic focal tank (24 cm diameter, 4 mm thickness) placed at the center of a larger square glass tank (54 × 54 × 31 cm; Figure 1B). Four rectangular display tanks (27 × 10 × 30 cm) were positioned inside the main tank, one on each side, at a distance of 4 cm from the focal tank. All joints were sealed using DOWSIL™ GP silicone sealant to prevent leakage. The main tank was elevated on a raised platform, and its outer walls were covered with opaque paper to block external visual cues. The adjoining sides of the display tanks were lined with laminated black paper to prevent visual interaction among display fish.

**Figure 1.**
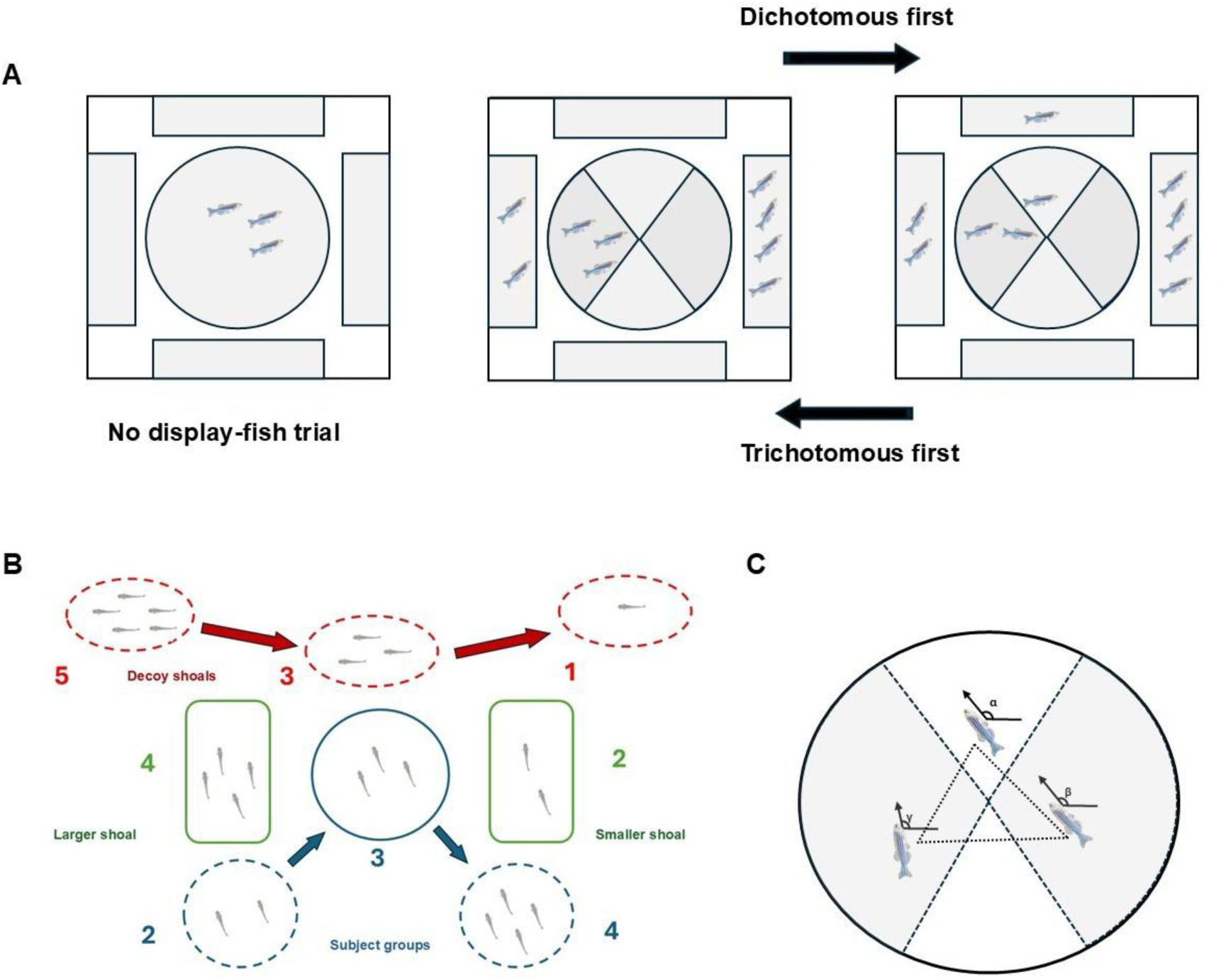
**A.** Schematic of the behavioral assay. In the no-display-fish trials, the subject group was placed in the central cylindrical arena with no stimulus shoals. In shoaling-choice trials, subjects experienced either a dichotomous presentation (larger vs. smaller shoal) or a trichotomous presentation (larger, smaller, and decoy shoal). Trial order was counterbalanced across subjects (dichotomous-first vs. trichotomous-first). **B.** Shoal size configurations for the dichotomous (larger vs. smaller) and trichotomous (larger, smaller, plus a decoy of size 1, 3, or 5) trials. Subject groups consisted of 2–4 fish (blue), and the target shoals were fixed at sizes 4 (larger) and 2 (smaller) (green). Decoys (red) appeared only in the trichotomous condition. **C.** Quantification of behavioral metrics in the focal arena. Inter-individual distance (IID) was calculated from pairwise distances between fish. Group polarization was derived from individual heading vectors (α, β, γ), and swimming speed was measured from frame-to-frame displacement. Stimulus shoals were positioned around a central cylindrical focal tank.

Each display tank was illuminated with an internal LED strip (Mufasa Copper Non-Waterproof LED Strip 5050) to eliminate shadows and maintain uniform lighting. Behavioral trials were recorded using a GoPro Hero 8 camera (1080p, 30 fps, linear mode) mounted on a tripod directly above the setup. To control for any potential bias caused by the presence of the tripod, an identical empty tripod was placed on the opposite side at the same height. The entire setup was enclosed within a 3 m × 3 m black photography cloth to minimize external visual distractions, ensuring that the only light sources were the display tanks.

The cylindrical design of the focal tank enabled the presentation of stimulus shoals from all four surrounding display tanks while reducing corner effects common in rectangular or square choice tanks. Although curved surfaces can introduce optical refraction, this was minimized by maintaining a water-filled space between the focal and display tanks. The rear walls of the display tanks were covered with green laminated sheets following the procedure described by Lucon-Xiccato et al. (2017).

### Experimental design

Male subjects in groups of 2, 3, or 4 fish were tested in a shoal-size preference task. Each group was presented with female display fish under two choice contexts: a dichotomous choice (4 vs. 2 fish) and a trichotomous choice (4 vs. 2 plus a decoy of 1, 3, or 5 fish). Each group was tested under three decoy conditions (decoy = 1, 3, or 5 fish) over three consecutive days, following one week of group housing in holding tanks within the fish system. Within each group-size and decoy condition, at least 20 groups were assigned to one presentation order (dichotomous-first or trichotomous-first), with the remaining groups assigned to the alternate order. The order of testing across group sizes was also randomized.

Before each trial, a cylindrical opaque plastic sheet was placed between the focal and display tanks to visually isolate the focal group from the display tanks. Each group first underwent a no-display fish trial, in which it was introduced into the central tank and visually isolated from the empty display tanks by a 2-minute period behind a cylindrical opaque plastic sheet. This control trial was used to assess any baseline spatial preferences as well as baseline group parameters, in the absence of display fish.

After the no-display trial, each group completed a shoaling preference task. Depending on the assigned order condition, subject groups experienced either the dichotomous-first condition, in which the binary choice (4 vs. 2 fish) preceded the trichotomous choice (4 vs. 2 plus a decoy of 1, 3, or 5 fish), or the trichotomous-first condition, in which the decoy trial was presented first. Each trial type (no-display, dichotomous, or trichotomous) lasted 3 minutes and was separated by a 2-minute period of visual isolation.

To minimize potential side biases, the display fish were randomly assigned to the four available display tanks. The axis of presentation, the side of the larger shoal, and the position of the decoy in trichotomous trials were also randomized.

### Fish tracking

The videos were processed using the multi-animal DeepLabCut (DLC), a Python-based pose estimation package (Lauer et al. 2022), to track the head, body, and tail of each fish in the group. A subset of videos was randomly selected to generate training frames, manually marking pre-defined body parts. The default Artificial Neural Network ResNet 50 was trained for more than 150,000 iterations. Labeled videos were generated for each group condition to verify tracking accuracy.

### Analysis

Trajectory data from each individual within each group were analyzed to derive four behavioral measures: (1) relative preference for the larger shoal, (2) mean inter-individual distance (IID), (3) mean polarization, and (4) mean individual speed. Group-level behavior was inferred using mixed-effects models, treating group ID as a random effect.

### Relative preference for the larger shoal

The circular arena was divided into two opposite angular sectors, one facing the larger shoal and the other facing the smaller shoal. Each 6 cm-wide sector, approximately twice the body length of an adult zebrafish, served as a preference zone for assessing shoaling behavior (Figure 1). Trajectory data were processed using the custom R script DLC-Analyzer (Sturman et al., 2020) to calculate the time spent by each individual of the group in each sector. The preference index (PI) for the larger shoal was computed as:

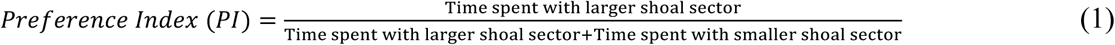

Preference for the larger shoal was modeled using a generalized linear mixed-effects model (GLMM), with the individual preference index as the response variable. Group size and the three-way interaction among decoy condition (1, 3, or 5), choice treatment (dichotomous or trichotomous), and order of presentation (dichotomous-first or trichotomous-first) were included as fixed effects. Group identity (group ID) was included as a random intercept to account for non-independence of individuals within the same group. The model structure was specified a priori based on the experimental design to minimize overfitting.

In addition to quantifying the decoy effect as the shift in relative preference between the dichotomous and trichotomous trials, we also calculated, for each group, the difference in relative preference between the two trial types (ΔPI). This difference was then modeled using a linear model (LM), with the preference difference as the response variable and group size and the interaction between decoy condition and order of presentation as fixed effects. Since each group produced a single value for the set of trials (the dichotomous and trichotomous conditions for that group), no random-effect structure was included.

### Time in zone

Time spent in each sector was analyzed using a generalized linear mixed-effects model (GLMM) with a beta error distribution, with the proportion of time spent in each zone as the response variable. Group size and a three-way interaction among zone ID, choice treatment (dichotomous or trichotomous), and order of presentation (dichotomous-first or trichotomous-first) were included as fixed effects. Separate models were fitted for each decoy conditions (1, 3, or 5). Group identity (group ID) was included as a random intercept to account for non-independence of individuals within the same group. The model structure was specified a priori based on the experimental design to minimize overfitting.

### Inter-individual Distances (IID)

For each group, the inter-individual distance (IID) at each time point was computed as the shortest distance between the body-axis line segments (joining the head and tail positions) of every pair of fish, using a custom Python script. This yielded one IID for groups of two fish, three IIDs for groups of three fish, and six IIDs for groups of four fish, corresponding to all pairwise combinations.

The mean IID for the duration of recording was then calculated for each individual and modeled using a linear mixed-effects model (LMM), with the individual mean IID as the response variable. Group size and the three-way interaction among decoy conditions (1, 3, or 5), choice treatment (dichotomous or trichotomous), and order of presentation (dichotomous-first or trichotomous-first) were included as fixed effects. Group identity (group ID) was included as a random intercept to account for non-independence.

### Group Polarization

Heading angles were computed from the head and tail coordinates of each fish using the arctan2 function in Python (arctan2(Δy, Δx)), where Δx and Δy represent the horizontal and vertical components of the body-axis vector (head minus tail). This yielded the orientation of each fish in radians, measured relative to the horizontal axis and ranging from −π to π.

The group polarization at each time point was then calculated as:

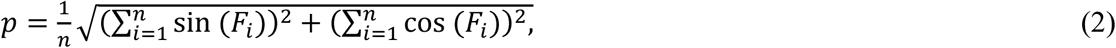

Where 𝐹_𝑖_ is heading angle of the fish 𝑖.

The mean polarization was calculated for each trial (dichotomous and trichotomous) for each group, and the data were modeled using a linear mixed-effects model (LMM), with mean polarization as the response variable.Group size and the three-way interaction among decoy condition (1, 3, or 5), choice treatment (dichotomous or trichotomous), and order of presentation (dichotomous-first or trichotomous-first) were included as fixed effects. Group identity (group ID) was included as a random intercept to account for non-independence.

### Speed

The mean speed for each individual within each group was calculated using the trajR package in R, which computes instantaneous speed between successive trajectory points (i.e., the speed of each step). The mean of these stepwise speeds over the duration of the recording was then obtained for each individual. These mean individual speeds were analyzed using a linear mixed-effects model (LMM), with individual mean speed as the response variable. Group size and the three-way interaction among decoy condition (1, 3, or 5), choice treatment (dichotomous or trichotomous), and order of presentation (dichotomous-first or trichotomous-first) were included as fixed effects. Group identity (group ID) was included as a random intercept to account for non-independence among individuals within the same group.

### No-display fish trials

The no-display fish data for each group were analyzed separately for each response variable (excluding relative preference) to characterize baseline group behavior across different group sizes. A linear mixed-effect model (LMM) with group size as a fixed effect and group ID as random intercept was used for this analysis.

## Results

### Relative preference for the larger shoal

In the dichotomous-first order, fish exhibited a baseline preference for the larger shoal in the dichotomous treatment across all group sizes among all decoy treatments. For decoy 1 and 5, dichotomous PIs showed strong preference for the larger shoal (**Figure 2A**, **Table 1**; decoy 1: 0.64, 95% CI: 0.59, 0.68; decoy 5: 0.63, 95% CI: 0.58, 0.68 at group size 2), whereas the corresponding trichotomous estimates remained at chance levels (**Figure 2A**, **Table 1**; decoy 1: 0.55, 95% CI: 0.50, 0.60; decoy 5: 0.50, 95% CI: 0.45, 0.55), thus introducing extreme decoys reduced relative preference to indifference. In contrast, decoy 3 produced nearly identical PIs under both treatments (**Figure 2A**, **Table 1**; 0.63, 95% CI: 0.57, 0.68 in dichotomous vs. 0.63, 95% CI: 0.57, 0.67 in trichotomous at group size 2). Hence decoy 3 did not show contextual altering of relative preference for the larger shoal.

**Figure 2.**
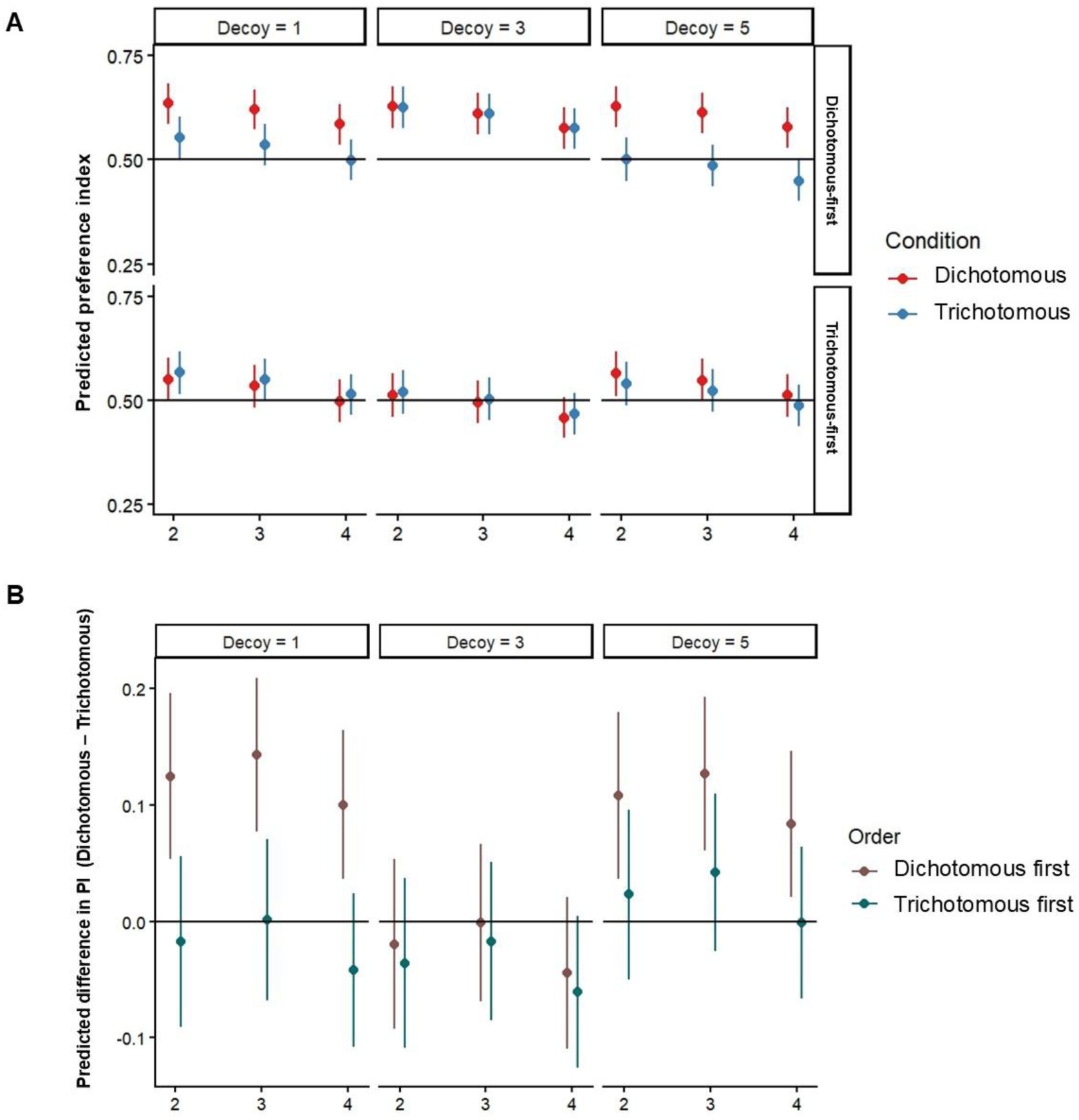
**A**. Model-estimated preference index (PI) for the larger shoal under dichotomous (red) and trichotomous (blue) conditions across group sizes (2, 3, 4) and decoy levels (1, 3, 5), with predicted means with 95% CIs. Panels indicate presentation order (dichotomous first and trichotomous first). The horizontal line at 0.5 indicates no preference for the larger shoal. **B**. Model-estimated difference in PI (ΔPI = dichotomous − trichotomous) across group sizes and decoy levels, with predicted means and 95% CIs. The horizontal line at 0 indicates no difference between dichotomous and trichotomous trials.

**Table 1:** GLMM estimates of the preference index (PI) with 95% confidence intervals for the effects of group size, decoy condition, choice treatment, and presentation order.

| Decoy | Group Size | Treatment | Estimate (PI) | 95% CI |
| --- | --- | --- | --- | --- |
| **Dichotomous-first** |  |  |  |  |
| 1 | 2 | Dicho | **0.64** | **0.59, 0.68** |
| 1 | 2 | Tricho | 0.55 | 0.50, 0.60 |
| 1 | 3 | Dicho | **0.62** | **0.57, 0.67** |
| 1 | 3 | Tricho | 0.54 | 0.49, 0.59 |
| 1 | 4 | Dicho | **0.59** | **0.54, 0.63** |
| 1 | 4 | Tricho | 0.50 | 0.45, 0.55 |
| 3 | 2 | Dicho | **0.63** | **0.57, 0.68** |
| 3 | 2 | Tricho | **0.63** | **0.57, 0.67** |
| 3 | 3 | Dicho | **0.61** | **0.56, 0.66** |
| 3 | 3 | Tricho | **0.61** | **0.56, 0.66** |
| 3 | 4 | Dicho | **0.58** | **0.52, 0.63** |
| 3 | 4 | Tricho | **0.58** | **0.52, 0.62** |
| 5 | 2 | Dicho | **0.63** | **0.58, 0.68** |
| 5 | 2 | Tricho | 0.50 | 0.45, 0.55 |
| 5 | 3 | Dicho | **0.61** | **0.56, 0.66** |
| 5 | 3 | Tricho | 0.49 | 0.43, 0.54 |
| 5 | 4 | Dicho | **0.58** | **0.53, 0.62** |
| 5 | 4 | Tricho | 0.45 | 0.40, 0.50 |
| **Trichotomous-first** |  |  |  |  |
| 1 | 2 | Dicho | 0.55 | 0.50, 0.60 |
| 1 | 2 | Tricho | **0.57** | **0.51, 0.62** |
| 1 | 3 | Dicho | 0.53 | 0.48, 0.59 |
| 1 | 3 | Tricho | 0.55 | 0.50, 0.60 |
| 1 | 4 | Dicho | 0.50 | 0.45, 0.55 |
| 1 | 4 | Tricho | 0.51 | 0.46, 0.56 |
| 3 | 2 | Dicho | 0.51 | 0.46, 0.56 |
| 3 | 2 | Tricho | 0.52 | 0.47, 0.57 |
| 3 | 3 | Dicho | 0.50 | 0.44, 0.55 |
| 3 | 3 | Tricho | 0.50 | 0.45, 0.55 |
| 3 | 4 | Dicho | 0.46 | 0.41, 0.51 |
| 3 | 4 | Tricho | 0.47 | 0.42, 0.52 |
| 5 | 2 | Dicho | **0.56** | **0.51, 0.62** |
| 5 | 2 | Tricho | 0.54 | 0.49, 0.59 |
| 5 | 3 | Dicho | 0.55 | 0.50, 0.60 |
| 5 | 3 | Tricho | 0.52 | 0.47, 0.57 |
| 5 | 4 | Dicho | 0.51 | 0.46, 0.56 |
| 5 | 4 | Tricho | 0.49 | 0.44, 0.54 |

Confidence intervals that did not include 0.5 were interpreted as indicating a preference significantly different from chance (in the frequentist sense) and are shown in bold.

In the trichotomous-first order, PIs were similar across treatments for all decoy types and group sizes, and consistently overlapped with chance (e.g., **Figure 2A**, **Table 1**; decoy 1: 0.53, 95% CI: 0.48, 0.59 in dichotomous vs. 0.55, 95% CI: 0.50, 0.60 in trichotomous at group size 3), reflecting a generally weaker baseline preference when fish first encountered a trichotomous choice.

Pairwise contrasts were consistent with the observed effects. In the dichotomous-first order, decoy 1 and decoy 5 showed significantly higher preference under dichotomous than trichotomous choice (**Table 2**; decoy 1: estimate = 0.349, p = 0.0007; decoy 5: estimate = 0.517, p < 0.0001), whereas decoy 3 showed no difference (**Table 2**; estimate = 0.004, p = 0.97). In the trichotomous-first order, contrast estimates were near zero for all decoys (all p > 0.35), indicating no treatment effect when trichotomous choice was experienced first.

**Table 2.** Pairwise comparisons of the preference index (PI) for dichotomous vs. trichotomous choice across decoy type and presentation order.

| Decoy | Order | Estimate (log odds) | SE | z-ratio | p value |
| --- | --- | --- | --- | --- | --- |
| 1 | Dicho-first | **0.3492** | 0.103 | 3.388 | **0.0007** |
| 3 | Dicho-first | 0.0041 | 0.110 | 0.038 | 0.9697 |
| 5 | Dicho-first | **0.5166** | 0.105 | 4.920 | **<.0001** |
| 1 | Tricho-first | -0.0642 | 0.106 | -0.605 | 0.5449 |
| 3 | Tricho-first | -0.0318 | 0.105 | -0.302 | 0.7625 |
| 5 | Tricho-first | 0.10032 | 0.108 | 0.930 | 0.3523 |

Estimates are averaged over group size and presented on the log-odds scale. p-values are Tukey-adjusted for multiple comparisons. Positive contrast estimates indicate a higher relative preference in the dichotomous condition compared with the trichotomous condition.

### Difference in relative preference between dichotomous and trichotomous choice (ΔPI)

In the dichotomous-first order, ΔPI values were clearly positive for Decoy 1 and Decoy 5 across all group sizes, with confidence intervals excluding zero. At group size 2, ΔPI was 0.12 (95% CI: 0.05,0.19, **Figure 2B**, **Table 3**) for Decoy 1 and 0.10 (95% CI: 0.03,0.17, **Figure 2B**, **Table 3**) for Decoy 5, indicating consistently higher relative preference for the larger shoal under dichotomous than trichotomous choice. In contrast, ΔPI estimates for Decoy 3 remained close to zero across group sizes (**Figure 2B**, **Table 3**; −0.03, 95% CI: −0.10, 0.04 at group size 2).

**Table 3.** GLMM estimates of the ΔPI (Dichotomous - Trichotomous) with 95% confidence intervals for the effects of group size, decoy condition, and presentation order.

| Decoy | Group Size | Estimate (PI) | 95% CI |
| --- | --- | --- | --- |
| **Dichotomous-first** |  |  |  |
| 1 | 2 | **0.12** | **0.05, 0.19** |
| 1 | 3 | **0.14** | **0.07, 0.20** |
| 1 | 4 | **0.09** | **0.03, 0.15** |
| 3 | 2 | -0.03 | -0.10, 0.04 |
| 3 | 3 | -0.01 | -0.07, 0.06 |
| 3 | 4 | -0.05 | -0.12, 0.01 |
| 5 | 2 | **0.10** | **0.03, 0.17** |
| 5 | 3 | **0.12** | **0.06, 0.18** |
| 5 | 4 | **0.08** | **0.01, 0.14** |
| **Trichotomous-first** |  |  |  |
| 1 | 2 | -0.03 | -0.10, 0.05 |
| 1 | 3 | -0.01 | -0.07, 0.06 |
| 1 | 4 | -0.05 | -0.11, 0.01 |
| 3 | 2 | -0.04 | -0.11, 0.03 |
| 3 | 3 | -0.03 | -0.09, 0.04 |
| 3 | 4 | **-0.07** | **-0.13, -0.01** |
| 5 | 2 | 0.01 | -0.06, 0.08 |
| 5 | 3 | 0.03 | -0.03, 0.10 |
| 5 | 4 | -0.01 | -0.07, 0.05 |

Confidence intervals that did not include 0 were interpreted as significantly different from zero (in the frequentist sense) and are shown in bold.

In the trichotomous-first order, ΔPI estimates remained close to zero for all decoys, with confidence intervals overlapping with zero across nearly in all decoy and group size combinations. At group size 2, ΔPI values were, −0.03 (95% CI: −0.10, 0.05, **Figure 2B**, **Table 3**) for Decoy 1, 0.01 (95% CI: −0.06, 0.08, **Figure 2B**, **Table 3**) for Decoy 5, and −0.04 (95% CI: −0.11, 0.03, **Figure 2B**, **Table 3**) for Decoy 3, indicating no difference between dichotomous and trichotomous choice when fish first encountered the trichotomous condition. The only exception was a small negative shift for Decoy 3 at group size 4 (**Figure 2B**, **Table 3**; −0.07, 95% CI: −0.13, –0.01), where relative preference for the larger shoal in the trichotomous treatment was higher.

Pairwise contrasts of ΔPI supported these patterns. In the dichotomous-first order, Decoy 1 elicited significantly larger ΔPI than Decoy 3 (**Table 4**; estimate = 0.145, p = 0.001), and Decoy 5 also differed significantly from Decoy 3 (**Table 4**; estimate = -0.128, p = 0.004), indicating that only these two decoys reliably altered relative preference. The contrast between Decoy 1 and Decoy 5 was non-significant (**Table 4**; estimate: 0.017, p = 0.90), consistent with their comparable influence on ΔPI. In the trichotomous-first order, all pairwise contrasts were close to zero and non-significant (all p > 0.32).

**Table 4.** Pairwise comparisons of the ΔPI (differences in preference index) among decoy types (1, 3, and 5) across presentation order.

| **Contrast** | **Order** | **Estimate** | **SE** | **t-ratio** | **p value** |
| --- | --- | --- | --- | --- | --- |
| Decoy1 - Decoy3 | Dicho-first | **0.1454** | **0.0404** | **3.602** | **0.001** |
| Decoy1 – Decoy5 | Dicho-first | 0.0173 | 0.0397 | 0.435 | 0.9010 |
| Decoy3 – Decoy5 | Dicho-first | **-0.1281** | **0.0401** | **-3.199** | **0.0040** |
| Decoy1 - Decoy3 | Tricho-first | 0.0184 | 0.0414 | 0.444 | 0.8968 |
| Decoy1 – Decoy5 | Tricho-first | -0.0402 | 0.0412 | -0.976 | 0.5924 |
| Decoy3 – Decoy5 | Tricho-first | -0.0586 | 0.0408 | -1.437 | 0.3222 |

Estimates are averaged over group size. p-values are Tukey-adjusted for multiple comparisons.

### Time in zone

In the dichotomous trial of the dichotomous-first order, fish consistently spent the greatest proportion of time with the larger shoal (zone 4) across all group sizes. Mean time spent near the larger shoal (**Figure 3**, **Table 5**; zone 4: ∼0.28–0.40) was substantially higher than time spent with the smaller shoal (**Figure 3**, **Table 5**; zone 2: ∼0.17–0.24) or empty zones (**Figure 3**, **Table 5**; zone 0: ∼0.12–0.19), which showed overlapping confidence intervals.

**Figure 3.**
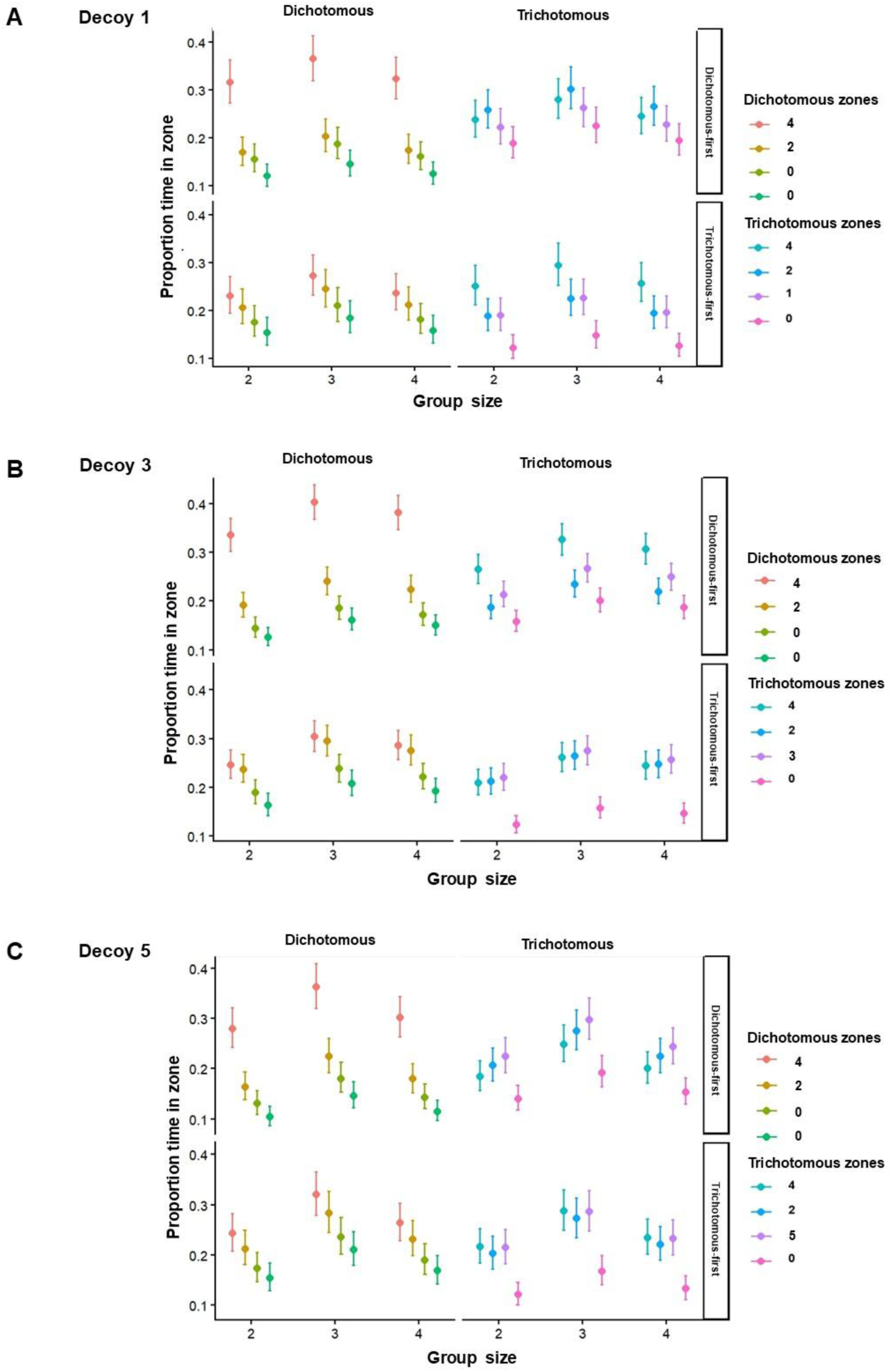
**A–C**. Model-estimated proportion of time spent in each zone under dichotomous and trichotomous conditions across group sizes (2, 3, 4) and decoy levels (**A**) 1, (**B**) 3, and (**C**) 5, with predicted means and 95% confidence intervals. Panels indicate presentation order (dichotomous first and trichotomous first).

**Table 5:** GLMM estimates of proportion of time in zones with 95% confidence intervals for the effects of group size, decoy condition, choice treatment, and presentation order.

| Group size | Order | Treatment | Zone | Estimate | 95% CI |
| --- | --- | --- | --- | --- | --- |
| Decoy 1 |  |  |  |  |  |
| 2 | Dicho-first | Dicho | 4 | 0.31 | 0.27, 0.36 |
| 2 | Dicho-first | Dicho | 2 | 0.17 | 0.14, 0.20 |
| 2 | Dicho-first | Dicho | 0 | 0.16 | 0.12, 0.18 |
| 2 | Dicho-first | Dicho | 0 | 0.12 | 0.09, 0.14 |
| 2 | Dicho-first | Tricho | 4 | 0.24 | 0.20, 0.27 |
| 2 | Dicho-first | Tricho | 2 | 0.26 | 0.21, 0.30 |
| 2 | Dicho-first | Tricho | 1 | 0.22 | 0.18, 0.26 |
| 2 | Dicho-first | Tricho | 0 | 0.19 | 0.15, 0.22 |
| 3 | Dicho-first | Dicho | 4 | 0.36 | 0.31, 0.41 |
| 3 | Dicho-first | Dicho | 2 | 0.20 | 0.17, 0.23 |
| 3 | Dicho-first | Dicho | 0 | 0.19 | 0.15, 0.22 |
| 3 | Dicho-first | Dicho | 0 | 0.15 | 0.12, 0.17 |
| 3 | Dicho-first | Tricho | 4 | 0.28 | 0.23, 0.32 |
| 3 | Dicho-first | Tricho | 2 | 0.30 | 0.25, 0.34 |
| 3 | Dicho-first | Tricho | 1 | 0.26 | 0.22, 0.30 |
| 3 | Dicho-first | Tricho | 0 | 0.22 | 0.18, 0.26 |
| 4 | Dicho-first | Dicho | 4 | 0.32 | 0.28, 0.36 |
| 4 | Dicho-first | Dicho | 2 | 0.17 | 0.14, 0.20 |
| 4 | Dicho-first | Dicho | 0 | 0.16 | 0.13, 0.19 |
| 4 | Dicho-first | Dicho | 0 | 0.12 | 0.10, 0.14 |
| 4 | Dicho-first | Tricho | 4 | 0.24 | 0.20, 0.28 |
| 4 | Dicho-first | Tricho | 2 | 0.26 | 0.22, 0.30 |
| 4 | Dicho-first | Tricho | 1 | 0.23 | 0.19, 0.26 |
| 4 | Dicho-first | Tricho | 0 | 0.19 | 0.16, 0.22 |
| 2 | Tricho-first | Dicho | 4 | 0.23 | 0.19, 0.27 |
| 2 | Tricho-first | Dicho | 2 | 0.21 | 0.17, 0.24 |
| 2 | Tricho-first | Dicho | 0 | 0.18 | 0.14, 0.21 |
| 2 | Tricho-first | Dicho | 0 | 0.15 | 0.12, 0.18 |
| 2 | Tricho-first | Tricho | 4 | 0.25 | 0.21, 0.29 |
| 2 | Tricho-first | Tricho | 2 | 0.19 | 0.15, 0.22 |
| 2 | Tricho-first | Tricho | 1 | 0.19 | 0.15, 0.22 |
| 2 | Tricho-first | Tricho | 0 | 0.12 | 0.10, 0.14 |
| 3 | Tricho-first | Dicho | 4 | 0.27 | 0.23, 0.31 |
| 3 | Tricho-first | Dicho | 2 | 0.25 | 0.20, 0.28 |
| 3 | Tricho-first | Dicho | 0 | 0.21 | 0.17, 0.24 |
| 3 | Tricho-first | Dicho | 0 | 0.19 | 0.15, 0.22 |
| 3 | Tricho-first | Tricho | 4 | 0.29 | 0.25, 0.34 |
| 3 | Tricho-first | Tricho | 2 | 0.23 | 0.19, 0.26 |
| 3 | Tricho-first | Tricho | 1 | 0.23 | 0.19, 0.26 |
| 3 | Tricho-first | Tricho | 0 | 0.15 | 0.12, 0.17 |
| 4 | Tricho-first | Dicho | 4 | 0.24 | 0.20, 0.27 |
| 4 | Tricho-first | Dicho | 2 | 0.21 | 0.17, 0.25 |
| 4 | Tricho-first | Dicho | 0 | 0.18 | 0.15, 0.21 |
| 4 | Tricho-first | Dicho | 0 | 0.16 | 0.13, 0.18 |
| 4 | Tricho-first | Tricho | 4 | 0.26 | 0.21, 0.29 |
| 4 | Tricho-first | Tricho | 2 | 0.19 | 0.16, 0.23 |
| 4 | Tricho-first | Tricho | 1 | 0.20 | 0.16, 0.23 |
| 4 | Tricho-first | Tricho | 0 | 0.13 | 0.10, 0.15 |
| Decoy 3 |  |  |  |  |  |
| 2 | Dicho-first | Dicho | 4 | 0.33 | 0.30, 0.36 |
| 2 | Dicho-first | Dicho | 2 | 0.19 | 0.16, 0.21 |
| 2 | Dicho-first | Dicho | 0 | 0.14 | 0.12, 0.16 |
| 2 | Dicho-first | Dicho | 0 | 0.13 | 0.10, 0.14 |
| 2 | Dicho-first | Tricho | 4 | 0.26 | 0.23, 0.29 |
| 2 | Dicho-first | Tricho | 2 | 0.19 | 0.16, 0.21 |
| 2 | Dicho-first | Tricho | 3 | 0.21 | 0.18, 0.24 |
| 2 | Dicho-first | Tricho | 0 | 0.16 | 0.13, 0.18 |
| 3 | Dicho-first | Dicho | 4 | 0.40 | 0.36, 0.43 |
| 3 | Dicho-first | Dicho | 2 | 0.24 | 0.21, 0.26 |
| 3 | Dicho-first | Dicho | 0 | 0.18 | 0.16, 0.20 |
| 3 | Dicho-first | Dicho | 0 | 0.16 | 0.14, 0.18 |
| 3 | Dicho-first | Tricho | 4 | 0.33 | 0.29, 0.35 |
| 3 | Dicho-first | Tricho | 2 | 0.23 | 0.20, 0.26 |
| 3 | Dicho-first | Tricho | 3 | 0.27 | 0.23, 0.29 |
| 3 | Dicho-first | Tricho | 0 | 0.20 | 0.17, 0.22 |
| 4 | Dicho-first | Dicho | 4 | 0.38 | 0.34, 0.41 |
| 4 | Dicho-first | Dicho | 2 | 0.22 | 0.19, 0.25 |
| 4 | Dicho-first | Dicho | 0 | 0.17 | 0.14, 0.19 |
| 4 | Dicho-first | Dicho | 0 | 0.15 | 0.12, 0.17 |
| 4 | Dicho-first | Tricho | 4 | 0.31 | 0.27, 0.33 |
| 4 | Dicho-first | Tricho | 2 | 0.22 | 0.19, 0.24 |
| 4 | Dicho-first | Tricho | 3 | 0.25 | 0.22, 0.27 |
| 4 | Dicho-first | Tricho | 0 | 0.19 | 0.16, 0.21 |
| 2 | Tricho-first | Dicho | 4 | 0.25 | 0.21, 0.27 |
| 2 | Tricho-first | Dicho | 2 | 0.24 | 0.20, 0.26 |
| 2 | Tricho-first | Dicho | 0 | 0.19 | 0.16, 0.21 |
| 2 | Tricho-first | Dicho | 0 | 0.16 | 0.14, 0.18 |
| 2 | Tricho-first | Tricho | 4 | 0.21 | 0.18, 0.23 |
| 2 | Tricho-first | Tricho | 2 | 0.21 | 0.18, 0.23 |
| 2 | Tricho-first | Tricho | 3 | 0.22 | 0.19, 0.24 |
| 2 | Tricho-first | Tricho | 0 | 0.12 | 0.10, 0.14 |
| 3 | Tricho-first | Dicho | 4 | 0.30 | 0.27, 0.33 |
| 3 | Tricho-first | Dicho | 2 | 0.29 | 0.26, 0.32 |
| 3 | Tricho-first | Dicho | 0 | 0.24 | 0.21, 0.26 |
| 3 | Tricho-first | Dicho | 0 | 0.21 | 0.18, 0.23 |
| 3 | Tricho-first | Tricho | 4 | 0.26 | 0.23, 0.29 |
| 3 | Tricho-first | Tricho | 2 | 0.26 | 0.23, 0.29 |
| 3 | Tricho-first | Tricho | 3 | 0.27 | 0.24, 0.30 |
| 3 | Tricho-first | Tricho | 0 | 0.16 | 0.13, 0.18 |
| 4 | Tricho-first | Dicho | 4 | 0.29 | 0.25, 0.31 |
| 4 | Tricho-first | Dicho | 2 | 0.28 | 0.24, 0.30 |
| 4 | Tricho-first | Dicho | 0 | 0.22 | 0.19, 0.24 |
| 4 | Tricho-first | Dicho | 0 | 0.19 | 0.16, 0.21 |
| 4 | Tricho-first | Tricho | 4 | 0.24 | 0.21, 0.27 |
| 4 | Tricho-first | Tricho | 2 | 0.25 | 0.21, 0.27 |
| 4 | Tricho-first | Tricho | 3 | 0.26 | 0.22, 0.28 |
| 4 | Tricho-first | Tricho | 0 | 0.15 | 0.12, 0.16 |
| Decoy 5 |  |  |  |  |  |
| 2 | Dicho-first | Dicho | 4 | 0.28 | 0.24, 0.32 |
| 2 | Dicho-first | Dicho | 2 | 0.16 | 0.13, 0.19 |
| 2 | Dicho-first | Dicho | 0 | 0.13 | 0.10, 0.15 |
| 2 | Dicho-first | Dicho | 0 | 0.10 | 0.08, 0.12 |
| 2 | Dicho-first | Tricho | 4 | 0.18 | 0.15, 0.21 |
| 2 | Dicho-first | Tricho | 2 | 0.20 | 0.17, 0.24 |
| 2 | Dicho-first | Tricho | 5 | 0.22 | 0.19, 0.26 |
| 2 | Dicho-first | Tricho | 0 | 0.14 | 0.11, 0.16 |
| 3 | Dicho-first | Dicho | 4 | 0.36 | 0.31, 0.40 |
| 3 | Dicho-first | Dicho | 2 | 0.22 | 0.19, 0.26 |
| 3 | Dicho-first | Dicho | 0 | 0.18 | 0.15, 0.21 |
| 3 | Dicho-first | Dicho | 0 | 0.14 | 0.12, 0.17 |
| 3 | Dicho-first | Tricho | 4 | 0.25 | 0.21, 0.28 |
| 3 | Dicho-first | Tricho | 2 | 0.27 | 0.23, 0.31 |
| 3 | Dicho-first | Tricho | 5 | 0.30 | 0.26, 0.34 |
| 3 | Dicho-first | Tricho | 0 | 0.20 | 0.16, 0.22 |
| 4 | Dicho-first | Dicho | 4 | 0.30 | 0.26, 0.34 |
| 4 | Dicho-first | Dicho | 2 | 0.18 | 0.15, 0.21 |
| 4 | Dicho-first | Dicho | 0 | 0.14 | 0.12, 0.17 |
| 4 | Dicho-first | Dicho | 0 | 0.11 | 0.09, 0.13 |
| 4 | Dicho-first | Tricho | 4 | 0.20 | 0.17, 0.23 |
| 4 | Dicho-first | Tricho | 2 | 0.22 | 0.19, 0.25 |
| 4 | Dicho-first | Tricho | 5 | 0.24 | 0.20, 0.28 |
| 4 | Dicho-first | Tricho | 0 | 0.15 | 0.12, 0.18 |
| 2 | Tricho-first | Dicho | 4 | 0.24 | 0.20, 0.28 |
| 2 | Tricho-first | Dicho | 2 | 0.21 | 0.18, 0.24 |
| 2 | Tricho-first | Dicho | 0 | 0.17 | 0.14, 0.20 |
| 2 | Tricho-first | Dicho | 0 | 0.15 | 0.12, 0.18 |
| 2 | Tricho-first | Tricho | 4 | 0.21 | 0.18, 0.25 |
| 2 | Tricho-first | Tricho | 2 | 0.20 | 0.17, 0.23 |
| 2 | Tricho-first | Tricho | 5 | 0.21 | 0.18, 0.25 |
| 2 | Tricho-first | Tricho | 0 | 0.12 | 0.10, 0.14 |
| 3 | Tricho-first | Dicho | 4 | 0.32 | 0.27, 0.36 |
| 3 | Tricho-first | Dicho | 2 | 0.29 | 0.24, 0.32 |
| 3 | Tricho-first | Dicho | 0 | 0.23 | 0.20, 0.27 |
| 3 | Tricho-first | Dicho | 0 | 0.21 | 0.17, 0.24 |
| 3 | Tricho-first | Tricho | 4 | 0.28 | 0.24, 0.32 |
| 3 | Tricho-first | Tricho | 2 | 0.27 | 0.23, 0.31 |
| 3 | Tricho-first | Tricho | 5 | 0.28 | 0.24, 0.32 |
| 3 | Tricho-first | Tricho | 0 | 0.16 | 0.14, 0.19 |
| 4 | Tricho-first | Dicho | 4 | 0.26 | 0.22, 0.30 |
| 4 | Tricho-first | Dicho | 2 | 0.23 | 0.19, 0.26 |
| 4 | Tricho-first | Dicho | 0 | 0.18 | 0.16, 0.22 |
| 4 | Tricho-first | Dicho | 0 | 0.16 | 0.14, 0.20 |
| 4 | Tricho-first | Tricho | 4 | 0.23 | 0.20, 0.27 |
| 4 | Tricho-first | Tricho | 2 | 0.22 | 0.18, 0.25 |
| 4 | Tricho-first | Tricho | 5 | 0.23 | 0.20, 0.26 |
| 4 | Tricho-first | Tricho | 0 | 0.13 | 0.11, 0.15 |

In the subsequent trichotomous trial of the dichotomous-first order, under Decoy 1 and Decoy 5, time spent with the larger shoal, smaller shoal, and decoy zones was broadly similar, with substantial overlap in 95% confidence intervals (e.g., Decoy 1, group size 3: zone 4 = 0.28, 95% CI: 0.23–0.32; zone 2 = 0.30, 95% CI: 0.25–0.34; decoy zone = 0.26, 95% CI: 0.22– 0.30, **Figure 3**, **Table 5**). In contrast, under Decoy 3, fish spent the greatest proportion of time with the larger shoal (e.g., group size 3: 0.33, 95% CI: 0.29–0.35, **Figure 3**, **Table 5**), followed by the decoy zone (**Figure 3**, **Table 5**; 0.27, 95% CI: 0.23–0.29) and then the smaller shoal (**Figure 3**, **Table 5**; 0.23, 95% CI: 0.20–0.26), preserving the relative preference for the larger shoal observed in the preceding dichotomous trial.

In the trichotomous trial of the trichotomous-first order, time spent across the larger shoal, smaller shoal, and decoy zones was more evenly distributed, particularly under Decoy 3 and Decoy 5, where mean proportions showed substantial overlap in their 95% confidence intervals (e.g., Decoy 5, group size 3: zone 4 = 0.28, 95% CI: 0.24–0.32; zone 2 = 0.27, 95% CI: 0.23–0.31; decoy zone = 0.28, 95% CI: 0.24–0.32, **Figure 3**, **Table 5**). In contrast, under Decoy 1, fish spent more time with the larger shoal (zone 4: ∼0.25–0.29) than with the smaller shoal or decoy zones (both ∼0.19–0.23, **Figure 3**, **Table 5**), although confidence intervals partially overlapped, suggesting modest differences in time allocation among zones.

In the subsequent dichotomous trial of the trichotomous-first order, time allocation showed a clearer ordering of zones for Decoy 1 and Decoy 5, with the largest proportion of time spent near the larger shoal (e.g., Decoy 1, group size 3: 0.27, 95% CI: 0.23–0.31, **Figure 3**, **Table 5**), followed by the smaller shoal (0.25, 95% CI: 0.20–0.28, **Figure 3**, **Table 5**), the zone where the decoy had previously been present (∼0.19–0.21, **Figure 3**, **Table 5**), and the empty zones (∼0.15–0.19, **Figure 3**, **Table 5**). In contrast, under Decoy 3, time spent with the larger and smaller shoals was comparable (e.g., group size 3: zone 4 = 0.30, 95% CI: 0.27–0.33; zone 2 = 0.29, 95% CI: 0.26–0.32, **Figure 3**, **Table 5**), followed by similar occupancy of the empty zones, indicating reduced distinction between the two shoal options.

### Inter-individual Distances (IID)

Groups of 3 and 4 showed consistently higher IID (cm) than groups of 2 in the no-fish trial (**Figure 4Ai**., **Table 6**; 2–3: estimate = –1.788, p < 0.01; 2–4: estimate = –1.685, p < 0.01). For the shoaling tests, there was a main effect of group size on mean IID (F (2, 123.1) = 36.81, p < 0.01), with larger groups consistently showing higher IID across both dichotomous and trichotomous trial types and across presentation orders. For example, for decoy 1 condition in the dichotomous trials of dichotomous-first order, mean IID increased from Group 2 (**Figure 4Aii., Table 7**; 5.82, 95% CI: 5.36, 6.28) to Group 3 (7.39, 95% CI: 6.96, 7.81) and Group 4 (8.09, 95% CI: 7.67, 8.50). Similarly, in the corresponding for decoy 1 condition dichotomous trials of the trichotomous-first order, mean IID increased from Group 2 (**Figure 4Aii., Table 7**; 6.58, 95% CI: 6.12, 7.03) to Group 3 (8.14, 95% CI: 7.72, 8.57) and Group 4 (8.84, 95% CI: 8.43, 9.26).

**Figure 4.**
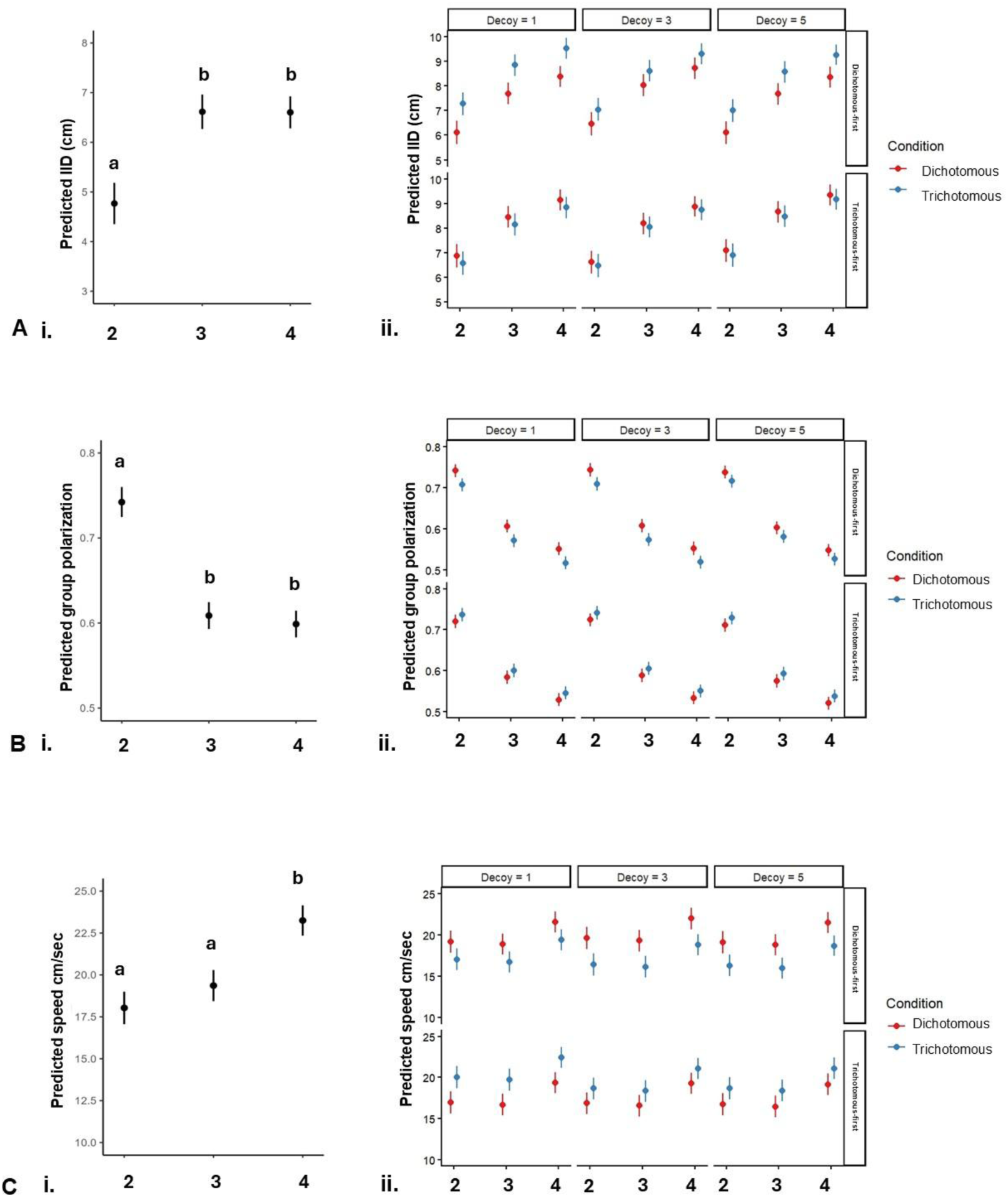
**A**. **Inter-individual distance (IID)**. Predicted IID for group sizes 2, 3, and 4 in trials **i**. without display fish and **ii**. for dichotomous and trichotomous choice sets across decoy types (1, 3, 5) and presentation-order conditions (dichotomous-first and trichotomous-first). **B**. **Group polarization**. Predicted group polarization for group sizes 2, 3, and 4 **i**. in trials without display fish and **ii**. for dichotomous and trichotomous choice sets across decoy types (1, 3, 5) and presentation-order conditions. **C**. **Speed**. Predicted swimming speed of individuals for group sizes 2, 3, and 4 **i**. in trials without display fish and **ii**. for dichotomous and trichotomous choice sets across decoy types (1, 3, 5) and presentation-order conditions. All values represent model-estimated marginal means ± 95% confidence intervals.

**Table 6:** Pairwise comparisons of IID, group polarization, and swimming speed across group sizes (2, 3, and 4) in no-display-fish trials.

| **Contrast** | **Estimate** | **SE** | **t-ratio** | **p value** |
| --- | --- | --- | --- | --- |
| Interindividual distance |  |  |  |  |
| Group 2 – Group 3 | **-1.788** | 0.267 | -6.707 | **<.0001** |
| Group 2 – Group 4 | **-1.685** | 0.262 | -6.422 | **<.0001** |
| Group 3– Group 4 | 0.104 | 0.242 | 0.427 | 0.9044 |
| Group polarisation |  |  |  |  |
| Group 2 – Group 3 | 0.1345 | 0.0108 | 12.408 | **<.0001** |
| Group 2 – Group 4 | 0.1446 | 0.0108 | 13.332 | **<.0001** |
| Group 3– Group 4 | 0.0101 | 0.0103 | 0.979 | 0.5914 |
| Speed |  |  |  |  |
| Group 2 – Group 3 | -1.33 | 0.644 | -2.067 | 0.1009 |
| Group 2 – Group 4 | **-5.23** | 0.635 | -8.238 | **<.0001** |
| Group 3– Group 4 | **-3.90** | 0.619 | -6.303 | **<.0001** |

p-values are Tukey-adjusted for multiple comparisons.

**Table 7:** GLMM estimates of the mean inter individual distances (IID in cm) with 95% confidence intervals for the effects of group size, decoy condition, choice treatment, and presentation order.

| Decoy | Group Size | Treatment | Order | Estimate (IID) | 95% CI |
| --- | --- | --- | --- | --- | --- |
| 1 | 2 | Dicho | DF | 5.82 | 5.36, 6.28 |
| 1 | 2 | Tricho | DF | 6.97 | 6.52, 7.42 |
| 1 | 3 | Dicho | DF | 7.39 | 6.96, 7.81 |
| 1 | 3 | Tricho | DF | 8.54 | 8.12, 8.96 |
| 1 | 4 | Dicho | DF | 8.09 | 7.67, 8.50 |
| 1 | 4 | Tricho | DF | 9.24 | 8.83, 9.65 |
| 3 | 2 | Dicho | DF | 6.16 | 5.70, 6.62 |
| 3 | 2 | Tricho | DF | 6.74 | 6.28, 7.19 |
| 3 | 3 | Dicho | DF | 7.73 | 7.30, 8.15 |
| 3 | 3 | Tricho | DF | 8.31 | 7.88, 8.73 |
| 3 | 4 | Dicho | DF | 8.43 | 8.01, 8.84 |
| 3 | 4 | Tricho | DF | 9.01 | 8.59, 9.42 |
| 5 | 2 | Dicho | DF | 5.80 | 5.35, 6.25 |
| 5 | 2 | Tricho | DF | 6.70 | 6.25, 7.15 |
| 5 | 3 | Dicho | DF | 7.37 | 6.95, 7.79 |
| 5 | 3 | Tricho | DF | 8.27 | 7.85, 8.68 |
| 5 | 4 | Dicho | DF | 8.07 | 7.66, 8.48 |
| 5 | 4 | Tricho | DF | 8.96 | 8.56, 9.37 |
| 1 | 2 | Dicho | TF | 6.58 | 6.12, 7.03 |
| 1 | 2 | Tricho | TF | 6.26 | 5.80, 6.72 |
| 1 | 3 | Dicho | TF | 8.14 | 7.72, 8.57 |
| 1 | 3 | Tricho | TF | 7.83 | 7.40, 8.26 |
| 1 | 4 | Dicho | TF | 8.84 | 8.43, 9.26 |
| 1 | 4 | Tricho | TF | 8.53 | 8.11, 8.95 |
| 3 | 2 | Dicho | TF | 6.32 | 5.86, 6.77 |
| 3 | 2 | Tricho | TF | 6.17 | 5.72, 6.63 |
| 3 | 3 | Dicho | TF | 7.88 | 7.46, 8.30 |
| 3 | 3 | Tricho | TF | 7.74 | 7.32, 8.16 |
| 3 | 4 | Dicho | TF | 8.58 | 8.18, 8.99 |
| 3 | 4 | Tricho | TF | 8.44 | 8.03, 8.85 |
| 5 | 2 | Dicho | TF | 6.78 | 6.33, 7.24 |
| 5 | 2 | Tricho | TF | 6.60 | 6.15, 7.06 |
| 5 | 3 | Dicho | TF | 8.35 | 7.93, 8.78 |
| 5 | 3 | Tricho | TF | 8.17 | 7.74, 8.60 |
| 5 | 4 | Dicho | TF | 9.05 | 8.64, 9.47 |
| 5 | 4 | Tricho | TF | 8.87 | 8.45, 9.29 |

A significant difference between IID in dichotomous versus trichotomous trials was observed only in the dichotomous-first order of presentation. Among the three group sizes, Group 3 showed the smallest difference (**Table 10**; estimate = –0.582, 95% CI: –0.900 to – 0.263, p < 0.01) compared to Group 2 (estimate = –1.152, 95% CI: –1.462 to –0.842, p < 0.01) and Group 4 (estimate = –0.896, 95% CI: –1.195 to –0.597, p < 0.01).

### Group polarization

Groups of 2 showed consistently higher group polarization than groups of 3 and 4 in the no-fish trial (**Figure 4Bi., Table 6**; 2–3: estimate = 0.1345, p < 0.01; 2–4: estimate = 0.1446, p < 0.01). For the shoaling tests, there was a main effect of group size on group polarization (χ² (2) = 751.84, p < 0.01), with larger groups consistently showing lower group polarization across both dichotomous and trichotomous trial types across presentation orders. For example, for decoy 1 condition in the dichotomous trials of the dichotomous-first order, decreased from Group 2 (**Figure 4Bii., Table 8**; 0.74, 95% CI: 0.73, 0.76) to Group 3 (0.61, 95% CI: 0.59, 0.62) and Group 4 (0.55, 95% CI: 0.54, 0.57). Similarly, in the corresponding for decoy 1 condition dichotomous trials of the trichotomous-first order, mean IID increased from Group 2 (**Figure 4Bii., Table 8**; 0.72, 95% CI: 0.71, 0.74) to Group 3 (0.59, 95% CI: 0.57, 0.60) and Group 4 (0.53, 95% CI: 0.52, 0.55).

**Table 8:** GLMM estimates of the group polarisation with 95% confidence intervals for the effects of group size, decoy condition, choice treatment, and presentation order.

| Decoy | Group Size | Treatment | Order | Estimate<br>(polarisation) | 95% CI |
| --- | --- | --- | --- | --- | --- |
| 1 | 2 | Dicho | DF | 0.74 | 0.73, 0.76 |
| 1 | 2 | Tricho | DF | 0.71 | 0.70, 0.73 |
| 1 | 3 | Dicho | DF | 0.61 | 0.59, 0.62 |
| 1 | 3 | Tricho | DF | 0.57 | 0.56, 0.59 |
| 1 | 4 | Dicho | DF | 0.55 | 0.54, 0.57 |
| 1 | 4 | Tricho | DF | 0.52 | 0.50, 0.53 |
| 3 | 2 | Dicho | DF | 0.75 | 0.73, 0.76 |
| 3 | 2 | Tricho | DF | 0.71 | 0.70, 0.73 |
| 3 | 3 | Dicho | DF | 0.61 | 0.59, 0.63 |
| 3 | 3 | Tricho | DF | 0.58 | 0.56, 0.59 |
| 3 | 4 | Dicho | DF | 0.56 | 0.54, 0.57 |
| 3 | 4 | Tricho | DF | 0.52 | 0.51, 0.54 |
| 5 | 2 | Dicho | DF | 0.74 | 0.73, 0.76 |
| 5 | 2 | Tricho | DF | 0.72 | 0.70, 0.73 |
| 5 | 3 | Dicho | DF | 0.61 | 0.59, 0.62 |
| 5 | 3 | Tricho | DF | 0.58 | 0.57, 0.60 |
| 5 | 4 | Dicho | DF | 0.55 | 0.54, 0.57 |
| 5 | 4 | Tricho | DF | 0.53 | 0.51, 0.54 |
| 1 | 2 | Dicho | TF | 0.72 | 0.71, 0.74 |
| 1 | 2 | Tricho | TF | 0.74 | 0.72, 0.76 |
| 1 | 3 | Dicho | TF | 0.59 | 0.57, 0.60 |
| 1 | 3 | Tricho | TF | 0.60 | 0.59, 0.62 |
| 1 | 4 | Dicho | TF | 0.53 | 0.52, 0.55 |
| 1 | 4 | Tricho | TF | 0.55 | 0.53, 0.56 |
| 3 | 2 | Dicho | TF | 0.73 | 0.71, 0.74 |
| 3 | 2 | Tricho | TF | 0.74 | 0.73, 0.76 |
| 3 | 3 | Dicho | TF | 0.59 | 0.58, 0.61 |
| 3 | 3 | Tricho | TF | 0.61 | 0.59, 0.62 |
| 3 | 4 | Dicho | TF | 0.54 | 0.52, 0.55 |
| 3 | 4 | Tricho | TF | 0.55 | 0.54, 0.57 |
| 5 | 2 | Dicho | TF | 0.71 | 0.70, 0.73 |
| 5 | 2 | Tricho | TF | 0.73 | 0.72, 0.75 |
| 5 | 3 | Dicho | TF | 0.58 | 0.56, 0.59 |
| 5 | 3 | Tricho | TF | 0.60 | 0.58, 0.61 |
| 5 | 4 | Dicho | TF | 0.52 | 0.51, 0.54 |
| 5 | 4 | Tricho | TF | 0.54 | 0.53, 0.56 |

**Table 9:** G LMM estimates of the mean speed (cm/sec) with 95% confidence intervals for the effects of group size, decoy condition, choice treatment, and presentation order.

| Decoy | Group Size | Treatment | Order | Estimate<br>(speed, cm/sec) | 95% CI |
| --- | --- | --- | --- | --- | --- |
| 1 | 2 | Dicho | DF | 19.03 | 17.77, 20.28 |
| 1 | 2 | Tricho | DF | 16.87 | 15.63, 18.11 |
| 1 | 3 | Dicho | DF | 18.72 | 17.50, 19.94 |
| 1 | 3 | Tricho | DF | 16.57 | 15.37, 17.77 |
| 1 | 4 | Dicho | DF | 21.42 | 20.21, 22.63 |
| 1 | 4 | Tricho | DF | 19.27 | 18.09, 20.45 |
| 3 | 2 | Dicho | DF | 19.46 | 18.18, 20.75 |
| 3 | 2 | Tricho | DF | 16.28 | 15.02, 17.54 |
| 3 | 3 | Dicho | DF | 19.16 | 17.91, 20.41 |
| 3 | 3 | Tricho | DF | 15.97 | 14.75, 17.20 |
| 3 | 4 | Dicho | DF | 21.86 | 20.62, 23.10 |
| 3 | 4 | Tricho | DF | 18.67 | 17.47, 19.88 |
| 5 | 2 | Dicho | DF | 18.96 | 17.71, 20.21 |
| 5 | 2 | Tricho | DF | 16.14 | 14.89, 17.38 |
| 5 | 3 | Dicho | DF | 18.66 | 17.45, 19.87 |
| 5 | 3 | Tricho | DF | 15.84 | 14.63, 17.04 |
| 5 | 4 | Dicho | DF | 21.35 | 20.17, 22.54 |
| 5 | 4 | Tricho | DF | 18.54 | 17.36, 19.71 |
| 1 | 2 | Dicho | TF | 16.84 | 15.56, 18.11 |
| 1 | 2 | Tricho | TF | 19.89 | 18.63, 21.16 |
| 1 | 3 | Dicho | TF | 16.53 | 15.30, 17.77 |
| 1 | 3 | Tricho | TF | 19.59 | 18.36, 20.83 |
| 1 | 4 | Dicho | TF | 19.23 | 18.01, 20.45 |
| 1 | 4 | Tricho | TF | 22.29 | 21.08, 23.50 |
| 3 | 2 | Dicho | TF | 16.73 | 15.47, 17.99 |
| 3 | 2 | Tricho | TF | 18.53 | 17.26, 19.79 |
| 3 | 3 | Dicho | TF | 16.43 | 15.20, 17.65 |
| 3 | 3 | Tricho | TF | 18.22 | 16.99, 19.46 |
| 3 | 4 | Dicho | TF | 19.12 | 17.92, 20.32 |
| 3 | 4 | Tricho | TF | 20.92 | 19.71, 22.13 |
| 5 | 2 | Dicho | TF | 16.63 | 15.36, 17.90 |
| 5 | 2 | Tricho | TF | 18.58 | 17.31, 19.86 |
| 5 | 3 | Dicho | TF | 16.33 | 15.09, 17.56 |
| 5 | 3 | Tricho | TF | 18.28 | 17.05, 19.52 |
| 5 | 4 | Dicho | TF | 19.03 | 17.81, 20.24 |
| 5 | 4 | Tricho | TF | 20.98 | 19.77, 22.20 |

**Table 10.** Pairwise comparisons of the IIDs for dichotomous vs trichotomous among decoy types (1, 3, and 5) across presentation order.

| Decoy | Order | Estimate | 95% CI | t-ratio | p value |
| --- | --- | --- | --- | --- | --- |
| 1 | Dicho-first | **-1.152** | -1.46, -0.84 | -7.290 | **<.0001** |
| 3 | Dicho-first | **-0.582** | -0.90, -0.26 | -3.581 | **0.0003** |
| 5 | Dicho-first | **-0.896** | -1.19, -0.59 | -5.879 | **<.0001** |
| 1 | Tricho-first | 0.313 | -0.006, 0.63 | 1.923 | 0.0545 |
| 3 | Tricho-first | 0.143 | -0.16, 0.44 | 0.916 | 0.3596 |
| 5 | Tricho-first | 0.181 | -0.13, 0.49 | 1.127 | 0.2601 |

Estimates are averaged over group size. p-values are Tukey-adjusted for multiple comparisons.

Group polarisation differed significantly between dichotomous and trichotomous trials across both presentation orders. A clear recency effect was observed, with polarisation being higher for the dichotomous trials in the dichotomous-first order and higher for the trichotomous trials in the trichotomous-first order for all group sizes (**Figure 4Bii., Table 8**, **Table 11)**.

**Table 11.**
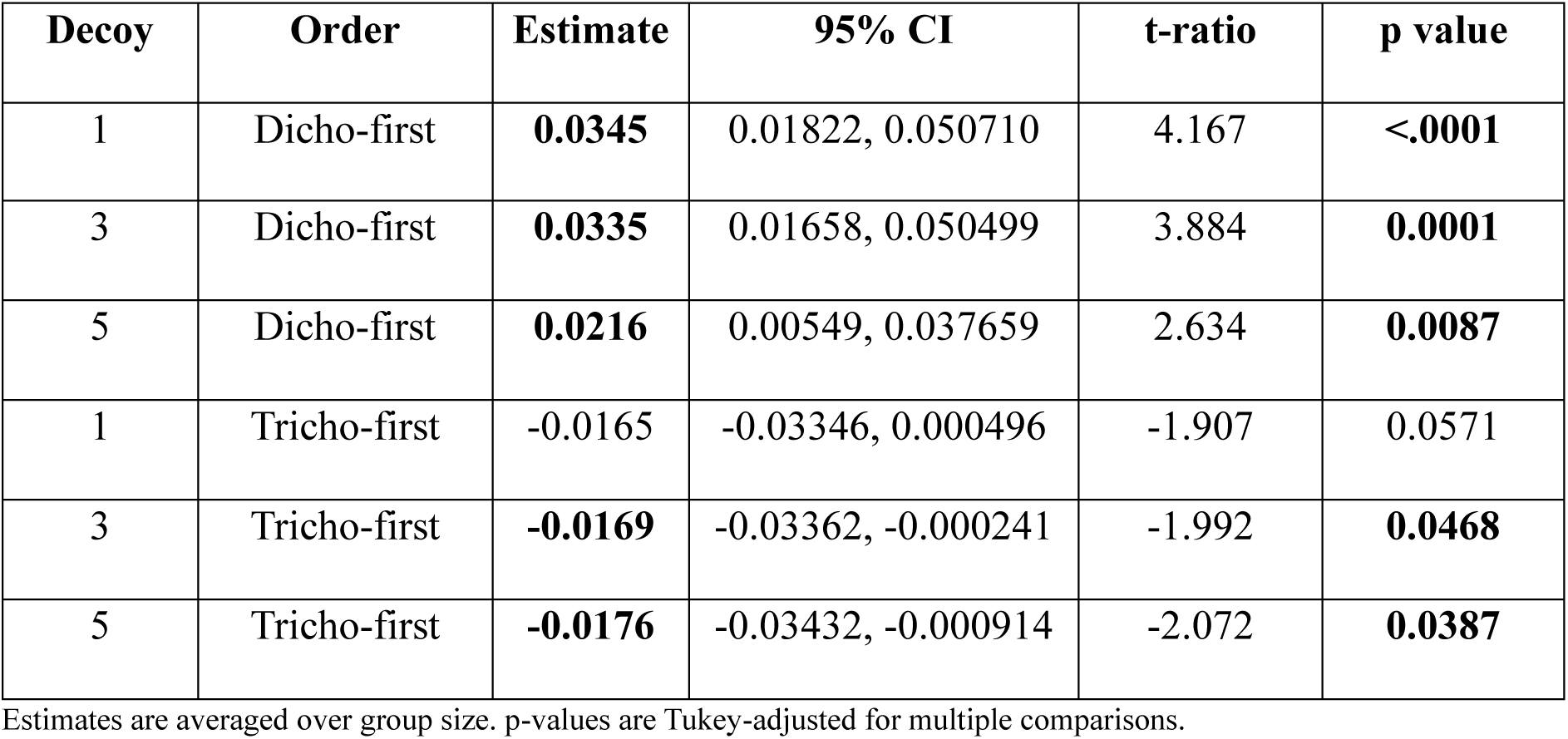
Pairwise comparisons of the group polarisation for dichotomous vs trichotomous among decoy types (1, 3, and 5) across presentation order.

| Decoy | Order | Estimate | 95% CI | t-ratio | p value |
| --- | --- | --- | --- | --- | --- |
| 1 | Dicho-first | **0.0345** | 0.01822, 0.050710 | 4.167 | **<.0001** |
| 3 | Dicho-first | **0.0335** | 0.01658, 0.050499 | 3.884 | **0.0001** |
| 5 | Dicho-first | **0.0216** | 0.00549, 0.037659 | 2.634 | **0.0087** |
| 1 | Tricho-first | -0.0165 | -0.03346, 0.000496 | -1.907 | 0.0571 |
| 3 | Tricho-first | **-0.0169** | -0.03362, -0.000241 | -1.992 | **0.0468** |
| 5 | Tricho-first | **-0.0176** | -0.03432, -0.000914 | -2.072 | **0.0387** |

Estimates are averaged over group size. p-values are Tukey-adjusted for multiple comparisons.

### Speed

Groups of 4 showed consistently higher mean speed (cm/sec) than groups of 2 and 3 in the no-fish trial (**Figure 4Ci., Table 6**; 2–4: estimate = -5.23, p < 0.01; 3–4: estimate = -3.90, p < 0.01). Similarly for the shoaling tests, the group of 4 showed higher speed compared to group size of 2 and 3 across both dichotomous and trichotomous trial types across presentation orders. For example, the mean speed in the decoy 1 condition in the dichotomous trials of the dichotomous-first order, decreased from Group 4 (**Figure 4Cii., Table 9**; 21.42, 95% CI: 20.21, 22.63) to Group 3 (18.72, 95% CI: 17.50, 19.94) and Group 2 (19.03, 95% CI: 17.77, 20.28).

Similarly, in the corresponding for decoy 1 condition dichotomous trials of the trichotomous-first order, mean speed increased from Group 4 (**Figure 4Cii., Table 9**; 19.23, 95% CI: 18.01, 20.45) to Group 3 (16.53, 95% CI: 15.30, 17.77) and Group 2 (16.84, 95% CI: 15.56, 18.11).

Mean speed differed significantly between dichotomous and trichotomous trials across both presentation orders. A clear recency effect was observed, with speed being higher for the dichotomous trials in the dichotomous-first order and higher for the trichotomous trials in the trichotomous-first order for all group sizes (**Figure 4Cii., Table 9**, **Table 12)**.

**Table 12.** Pairwise comparisons of mean speed for dichotomous vs trichotomous among decoy types (1, 3, and 5) across presentation order.

| **Decoy** | **Order** | **Estimate** | **SE** | **t-ratio** | **p value** |
| --- | --- | --- | --- | --- | --- |
| 1 | Dicho-first | **2.15** | 1.05, 3.256 | 3.833 | **0.0001** |
| 3 | Dicho-first | **3.19** | 2.04, 4.340 | 5.433 | **<.0001** |
| 5 | Dicho-first | **2.82** | 1.73, 3.905 | 5.089 | **<.0001** |
| 1 | Tricho-first | **-3.06** | -4.20, -1.918 | -5.259 | **<.0001** |
| 3 | Tricho-first | **-1.80** | -2.93, -0.664 | -3.110 | **0.0019** |
| 5 | Tricho-first | **-1.96** | -3.09, -0.817 | -3.369 | **0.0008** |

Estimates are averaged over group size. p-values are Tukey-adjusted for multiple comparisons.

## Discussion

While group decision-making is often proposed to buffer against individual biases. Whether increasing group size attenuates context-dependent economic biases remains untested. Our findings reveal that group size doesn’t influence shoal size preference and decoy-induced biases. The contextual effect of presenting a decoy shoal emerged only when the subject shoal first experienced a simple (dichotomous) choice first followed by the introduction of decoy shoal, and only when the decoys were extreme relative to the target options. When subject shoals first encountered a more complex (trichotomous) choice, the relative preference for the larger shoal remained close to chance. Together with patterns in zone occupancy and group kinematics, these results suggest that group-level social preference in zebrafish is sensitive to contextual structure and recent experience, but only under specific presentation orders.

### Group shoal size preference and context effects

Contrary to our expectations, subject group size did not modulate shoal size preference either in the baseline dichotomous contrast or following the introduction of a decoy shoal. To our knowledge, this is the first study to examine how the size of the decision-making group influences preference for conspecific shoals of different sizes. One possibility is that the relatively small group sizes in our task (n = 2-4) were insufficient to generate detectable collective benefits. Along similar lines, Ward et al. (2008) predicted no improvement in decision accuracy at small group sizes, arguing that gains in accuracy arise only after several independent decisions accumulate; consequently, only larger groups (n ≥ 8) benefit sufficiently from social information.

We did not observe context effects in the group assay that resembled those detected in the single-subject task (Singh et al. 2022). In groups, adult male zebrafish consistently showed a strong baseline preference for the larger shoal across all group sizes, and context effects were expressed primarily as a loss of relative preference under the decoy 1 and decoy 5 conditions. Although the decoy 3 and decoy 5 conditions were not tested in the single-animal task, we did not observe, at the group level, the increase in relative preference for the larger shoal (above chance) in the dichotomous contrast following prior exposure to a trichotomous choice, an effect previously observed in single males with decoy 1, using the same setup and design. These differences suggest that group-level shoaling decisions can diverge markedly from individual-level shoaling preference behaviour. This highlights the value of studying behaviour under ecologically relevant social conditions and highlights the limited ecological validity of individual-based laboratory assays (Santos et al. 2021; Kondrakiewicz et al. 2019).

This lack of contextual shift in relative preference in the decoy 3 condition indicates that zebrafish do not indiscriminately integrate additional social stimuli but rather show selective susceptibility to contextual restructuring of choice options. The fact that only the extreme decoys (relative to target options) could alter discriminability suggests that a decoy must substantially alter the perceived relative value of the target options to affect choice. Computational models based on adaptive gain control argue that distractor stimuli can reshape subjective value differences through normalization-like mechanisms, thereby weakening preferences when distractors lie at the extremes of the feature space (Cheadle et al. 2014; Li et al. 2018). The extreme decoys expand the representational range over which values are encoded, compressing the effective difference between focal options and driving preferences toward indifference. In contrast, decoys that fall near the midpoint of the existing choice set exert minimal gain modulation and thus preserve baseline preferences.

Patterns of time-in-zone shed light on the ordering of the shoal options underlying the observed effects in relative preference. In the dichotomous-first order, where we detected a decoy effect, time allocation during the trichotomous trial mirrored the disrupted preference pattern: fish shoal spent comparable time with the 4-fish and 2-fish shoals, indicating indifference, while time spent with the decoy scaled with its shoal size - lowest for decoy 1 and highest for decoy 5. Under the decoy 3 condition, where no decoy effect was observed, fish shoal exhibited the most “rational” ordering of time allocation, spending the most time with the 4-fish shoal, followed by the 3-fish decoy and then the 2-fish shoal. In the trichotomous-first order, where no decoy effects emerged overall, zone occupancy during the trichotomous trial showed distinct patterns across decoy types. Under decoy 1, fish spent the most time with the 4 fish shoal and nearly equal time with the 2 fish and 1 fish shoals, consistent with a similarity-effect pattern in which the decoy competes with one primary option (Frederick et al. 2014). In contrast, under decoy 3 and decoy 5, fish distributed time almost equally across all three shoals, suggesting limited discrimination among options when presented simultaneously. Reflecting potential increased cognitive or perceptual load associated with tracking multiple large-sized shoals, hence reducing discriminability. The removal of the decoy in the subsequent dichotomous trial restored a size-based pattern of time allocation: most time at the larger shoal, then the smaller shoal, and least in the former decoy zone. These results indicate that the apparent similarity in shifts of relative preference for the larger shoal under decoy 1 and decoy 5 may arise from distinct underlying mechanisms, likely involving an interaction between more than once cognitive processes like between how shoal sizes are psychophysically scaled and the cognitive demands of tracking and discriminating groups of different size (Spitmaan et al. 2019).

The order dependent contextual effects may be driven by two complementary mechanisms. First, anchoring effects may operate, whereby the first decision context establishes the scale or strategy used for subsequent comparisons. Encountering two options first may prime a simpler discrimination rule, allowing the third stimulus to be more easily placed on that scale. Conversely, encountering three options first may fail to establish a simpler reference scale, reducing the contrast between options. Second, cognitive load or attention allocation may differ between orders. Trichotomous-first fish must initially parse and compare three social stimuli simultaneously, which may weaken preferences due to reduced attentional focus or higher sampling costs.

The order-dependent context effects, such as contrast and anchoring, have been documented in other non-human animals. The colonies of *Temnothorax* ants evaluate mediocre nests more favourably when first exposed to poorer alternatives (Doering et al. 2023). Similarly, in bumblebees the relative attractiveness of medium sucrose-concentration flowers depended on presentation order: when medium flowers were first contrasted with high-concentration flowers, they were relatively inferior, but after a low-concentration alternative was introduced the medium option became comparatively more favourable (Hemingway et al. 2024). This shift has been attributed to an incentive contrast effect, a sensory mechanism in which prior exposure to low or high sucrose concentrations modulates bees’ perceptual thresholds (Bitterman 1976, Wiegmann et al. 2009) and which has been reported in bees as facilitating economic choice behaviour (Waldron et al. 2005). Apart from the incentive contrast effects, the loss of relative preference between two targets following the introduction of a third option has also been explained by the random dilution effect, wherein choice behaviour contains a stochastic component (Bateson et al. 2002). When a third option is introduced, it absorbs a fraction of these undirected or noise-driven choices. Although the underlying preference between the two primary options remains unchanged, the redistribution of stochastic selections reduces their apparent preference difference, revealing a weakened or flattened preference rather than a genuine shift in relative value (Bateson et al. 2002). We could control for contrast effects by inferring from the baseline preference from the dichotomous-first and trichotomous-first order conditions; however, the possibility that the observed shift in relative preference was driven by random dilution could not be excluded in our design.

### Group kinematics and shoal size preference

Group size systematically influenced group kinematics across both no-display and shoaling preference trials. In the no-fish trials, inter-individual distances (IID) increased in larger groups of 3 and 4 compared to the groups of 2. Although a similar trend in the influence of group size on IID has been reported in zebrafish (Suriyampola et al. 2022), these patterns may be driven by density rather than group size alone (Shelton et al. 2015), as groups of 4 are effectively more densely packed within the same physical arena than groups of 2. Group polarization showed the opposite pattern, with groups of 2 exhibiting the highest alignment and larger groups showing diminished coordination. This is also consistent from previous studies showing that larger zebrafish shoals are less polarized than smaller shoals (Miller & Gerlai 2012). Suriyampola et al. (2022) also show that in larger groups, individual group members spent less time in the leading position, suggesting more diffuse leadership. This shift in leader-follower structure could contribute to the less polarization observed among the larger groups in our study. Swim speed also differed by group size: groups of 4 swam significantly faster than groups of 2 and 3, whereas groups of 2 and 3 did not differ. Similar trends have been reported in other cyprinids, with swim speed rising as group size increases (Fu 2016), possibly due to reduced vigilance and the onset of schooling-like behavior in larger groups (Fu 2016; Miller & Gerlai 2012).

In the shoal size preference test, group size exerted a more linear influence on IID than in the no-fish trials, with IID progressively increasing from groups of 2 to 3 to 4. Of the kinematic measures examined, IID covaried most consistently with the patterns of relative preference for the larger shoal. IID was significantly higher in the trichotomous than in the dichotomous condition when the dichotomous task was presented first, indicating greater spacing in the presence of the decoy. This effect was weaker in the Decoy-3 condition, where no decoy effect was detected in relative preference and absent in the trichotomous-first order, which again mirrored the preference data. By contrast, IID patterns did not generalize to polarization or speed. Polarization declined linearly with increasing group size, and dichotomous shoals were marginally more polarized than trichotomous shoals only in the dichotomous-first condition. Speed showed a distinct pattern: groups of 2 and 3 swam at comparable speeds, whereas groups of 4 were consistently faster, similar to the no-fish trials. However, speed showed a task-order recency effect, with dichotomous-first shoals swimming faster in the dichotomous task and trichotomous-first shoals swimming faster in the trichotomous task.

Group polarization is often highlighted as a key metric mediating collective motion and decision-making (MacGregor et al. 2020; Davis et al. 2017) and frequently correlated with inter-individual spacing and speed (Kotrschal et al. 2020). In our study, however, the correspondence between IID and relative preference, but not with polarization, is notable. IID tracked the strength of the group’s preference for the larger shoal, suggesting that in this spatially distributed shoal choice task and at smaller group sizes, inter-individual spacing may be a more sensitive indicator of collective decision dynamics than polarization. This may reflect the weaker alignment pressures in small shoals, where the group’s collective motion is less constrained than in larger, highly coordinated schools.

## Conclusion

Our findings demonstrate that collective shoal size preference in zebrafish is influenced more strongly by the structure and order of shoal choice presentation than by the size of the decision-making group. Groups consistently preferred the larger shoal under baseline dichotomous choice, but the introduction of decoy shoals selectively disrupted relative preference, particularly when decoys were extreme and when a simpler dichotomous contrast preceded the trichotomous trial. The collective shoal preference is sensitive to recent experience and contextual restructuring of alternatives, and that group-level processes do not buffer or eliminate context-dependent biases.

The behavioural mechanisms underlying these effects likely include both psychophysical and collective components. Extreme decoys may alter perceived value differences between options via normalization-like processes, compressing discriminability and reducing relative preference, whereas intermediate decoys exert minimal influence. These shifts in preference were mirrored in inter-individual distance (IID), which increased under conditions in which preferences collapsed toward indifference, suggesting more distributed sampling or attentional splitting across options, whereas tighter spacing corresponded to stronger consensus and clearer preference. IID therefore provided a more sensitive correlate of decision state than polarization or swim speed in this spatially distributed task and at the small group sizes tested.

Together, these results indicate that collective shoal preference emerges from interactions among contextual structure, cognitive constraints, and spatial organization, rather than arising exclusively from group size or the simple aggregation of individual-level biases. This work underscores the importance of studying shoaling decision-making under ecologically relevant social conditions and cautions against extrapolating individual-based laboratory results to group-level behaviour.

